## Supplementary File2 for "A structured evaluation of genome-scale constraint-based modeling tools for microbial consortia"

**Table S1. Summarized comparison of the available** **static tools/approaches.**

| Modeling Tool/approach (Year Developed) | Website/GitHub Link | Optimization Routine (Single or Bilevel) | | Programming Language | Environment Dependencies | Optimization Dependencies | Namespace Requirement | # of citations (as of December 2022) |
| --- | --- | --- | --- | --- | --- | --- | --- | --- |
| OptCom (2012) | http://www.maranasgroup.com/submission/OptCom.htm | Bilevel | GAMS | |  | BARON | Yes | 285 |
| cFBA (2013) | https://journals.plos.org/plosone/article?id=10.1371/journal.pone.0064567 | Single | Python | | CBMPy | CPLEX | No | 149 |
| CASINO  (2015) | Not Available | Bilevel | MATLAB | | RAVEN | No information | No | 271 |
| SteadyCom  (2017) | https://github.com/opencobra/cobratoolbox/tree/master/src/analysis/multiSpecies/SteadyCom | Single | MATLAB | | COBRA Toolbox | LP solver and CPLEX | No | 133 |
| RedCom  (2019) | Not Available | Bilevel | MATLAB | | *CellNetAnalyzer* | CPLEX, *efmtool, fmincon* | No | 24 |
| MICOM  (2020) | https://micom-dev.github.io/micom/ | Single | Python | | COBRApy | OSQP  or CPLEX, Gurobi | No | 39 |
| MMT (v1, 2018; v2,2022) | https://github.com/opencobra/cobratoolbox/blob/master/src/analysis/multiSpecies/microbiomeModelingToolbox/README.md | Single | MATLAB | | COBRA Toolbox | LP solver | No | 79+3 |
| NECom  (2020) | <https://github.com/Jingyi-Cai/NECom.git>. | Bilevel | MATLAB | |  | BARON and Gurobi, CPLEX and glpk | No | 12 |

**Table** **S2. Rubric - Qualitative assessment of tools/approaches.**

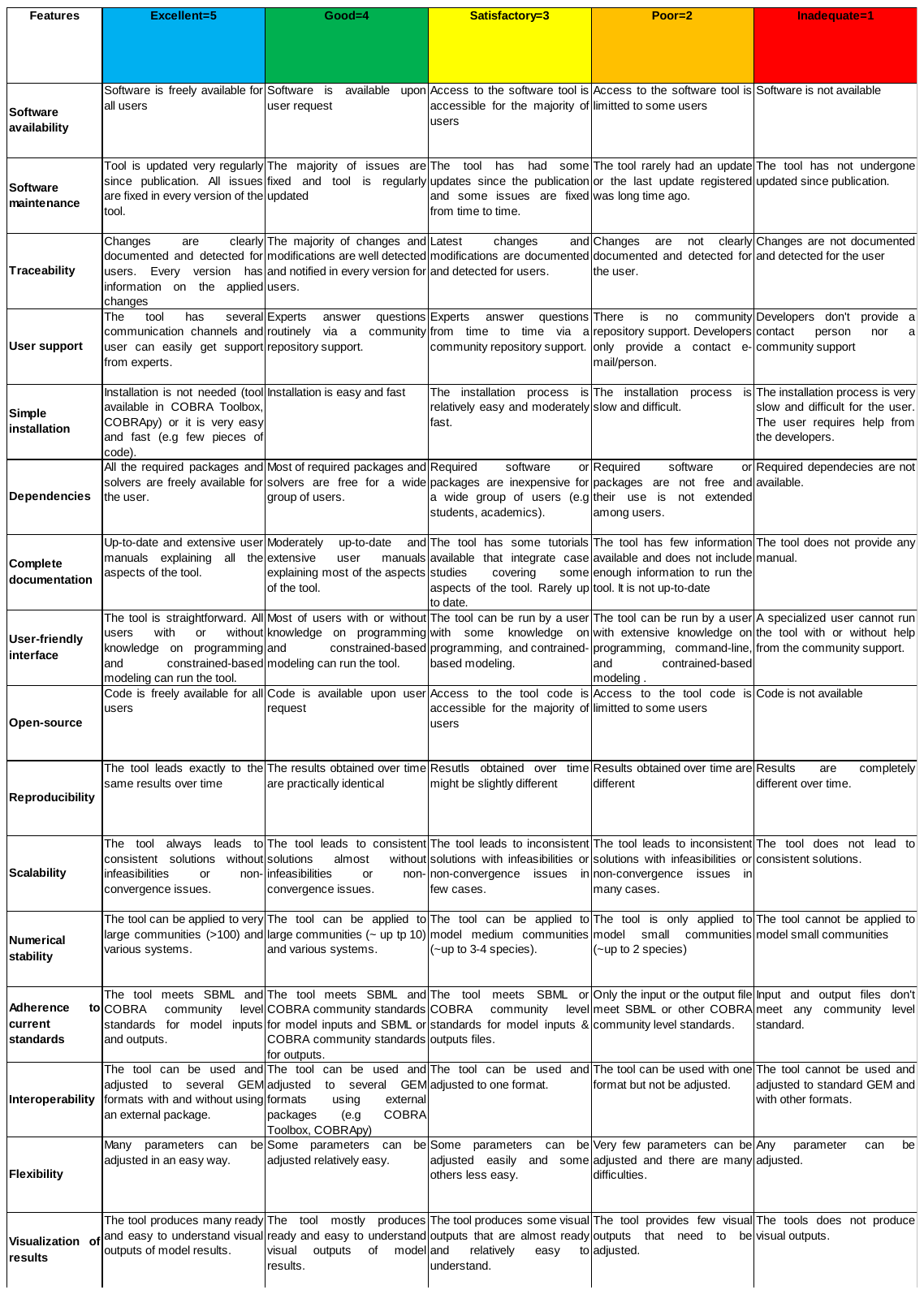

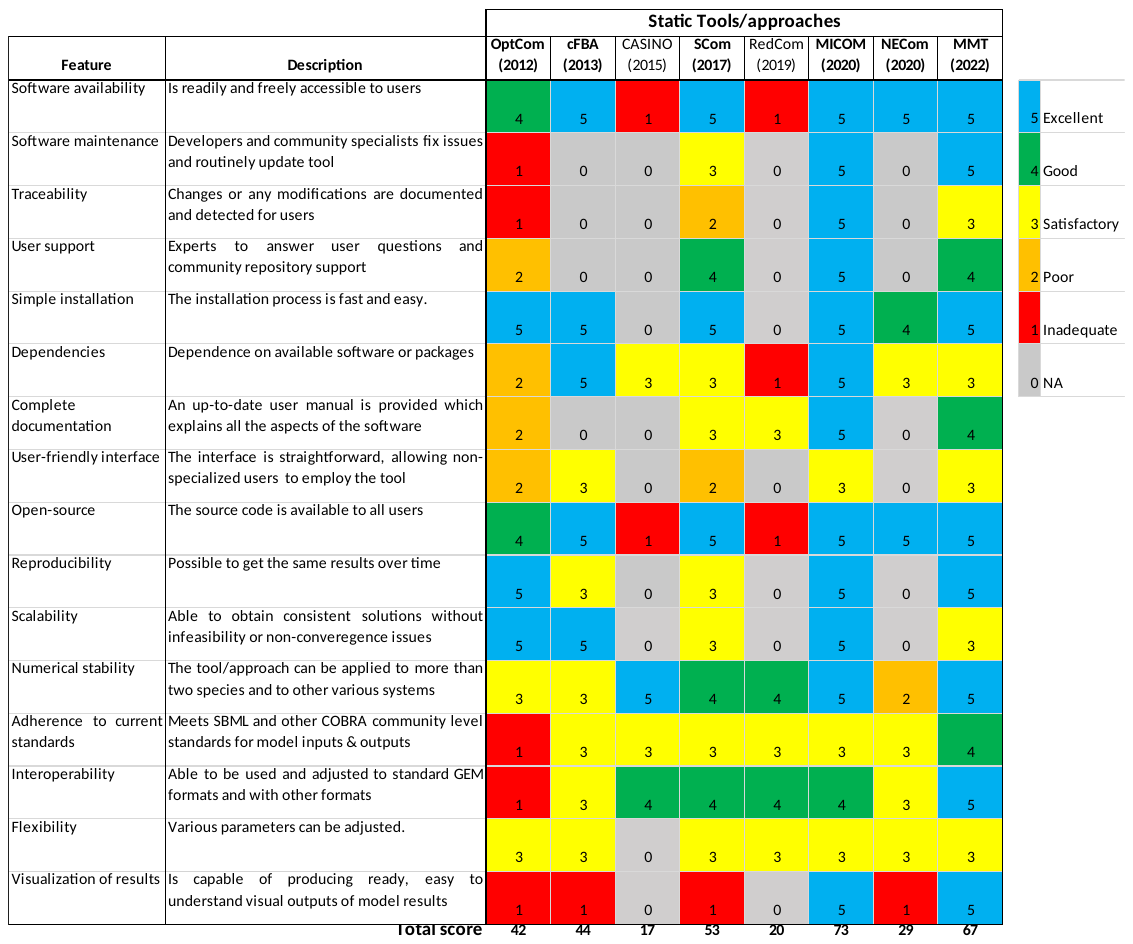

**Figure** **S1. Qualitative assessment of the static tools/approaches. Colored squared indicate the evaluation of the specified feature in every tool/approach. The color scale (upper right) goes from excellent (blue=5) to inadequate (red=1). When a feature does not apply to the specified tool/approach or the feature was not evaluated, it is indicated as NA (Not applicable; grey=0).The tools/approaches are ordered by year of publication. Scom: SteadyCom; MMT: Microbiome Modelling Toolbox.**

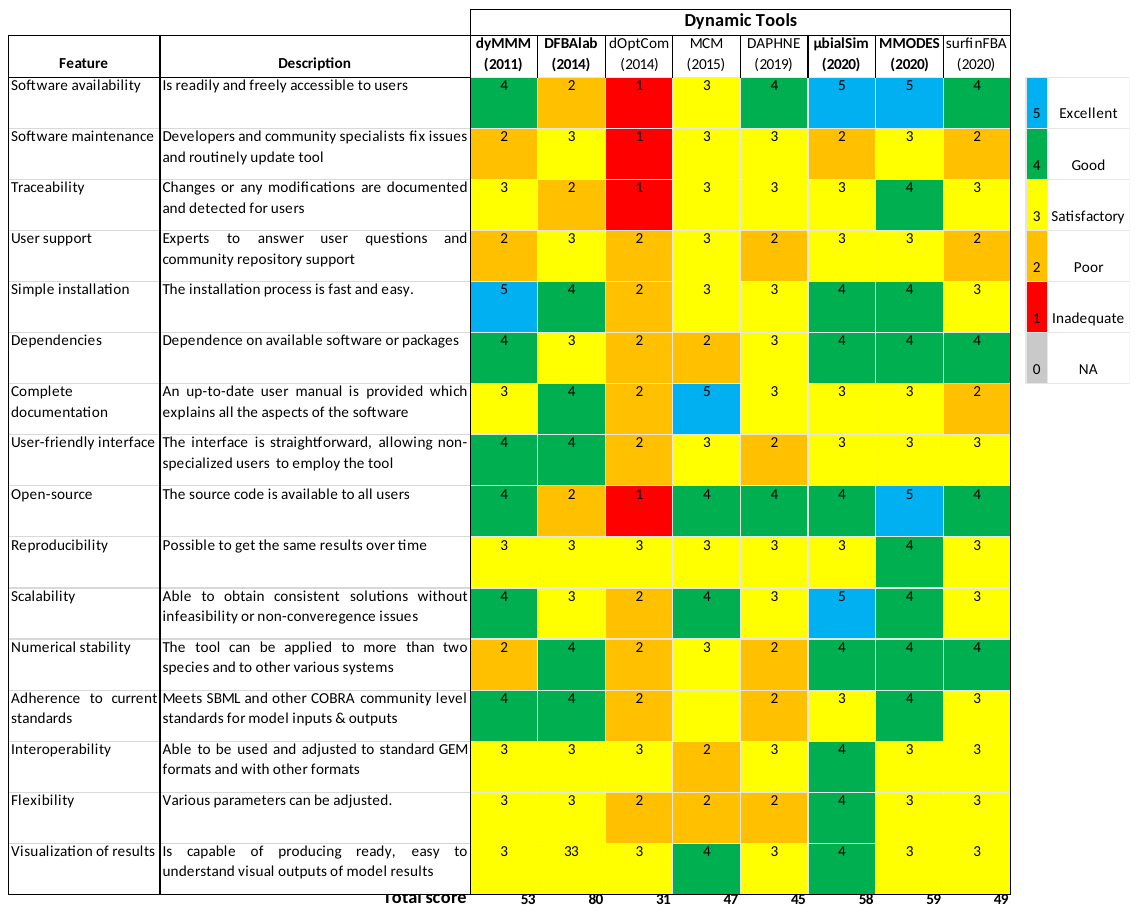

**Figure** **S2. Qualitative assessment of the dynamic tools.** Colored squared indicate the evaluation of the specified feature in every tool. The color scale (upper right) goes from excellent (blue) to inadequate (red). When a feature does not apply to the specified tool or the feature was not evaluated, it is indicated as NA (Not applicable; grey). The tools are ordered by year of publication.

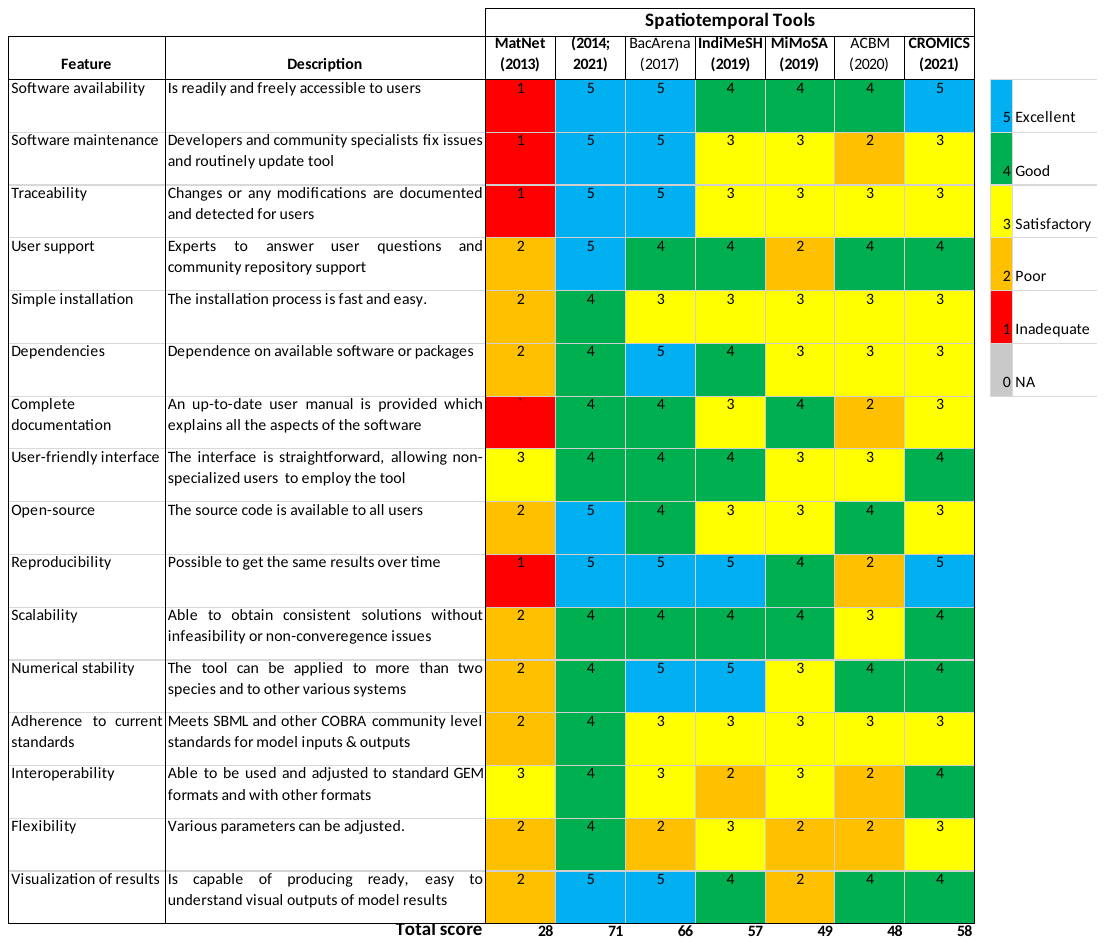

**Figure** **S3. Qualitative assessment of the spatiotemporal tools.** Colored squared indicate the evaluation of the specified feature in every tool. The color scale (upper right) goes from excellent (blue) to inadequate (red). When a feature does not apply to the specified tool or the feature was not evaluated, it is indicated as NA (Not applicable; grey). The tools are ordered by year of publication.

**Table** **S3**. **Inputs, outputs, and assumptions of the static tools/approaches.**

| Tool/approach | Inputs | Outputs | Assumptions |
| --- | --- | --- | --- |
| OptCom | GEM of single species, CO uptake rate, relative abundance, min. growth rate of species, total biomass. | Community growth rate, species growth rate, fluxes | Steady-state, species don’t grow at balanced growth. |
| cFBA | Community GEM, CO uptake rate, relative abundance, total biomass, species growth rate. | Community growth rate, fluxes | Steady-state, equal growth rates of species and community. |
| SteadyCom | GEM of single species, CO uptake rate, total biomass. | Relative abundance, Community growth rate, species growth rate, fluxes | Steady-state, equal growth rates of species and community. |
| MMT | GEM of single species, CO uptake rate, relative abundance,  total biomass, species growth rate. | Community growth rate, fluxes | Steady-state, equal growth rates of species and community. |
| MICOM | GEM of single species, CO uptake rate, relative abundance, total biomass. | Community growth rate, species growth rate, fluxes | Steady-state, species don’t grow at balanced growth. |

**Table S4**. **Genome-scale metabolic models (GEMs) and input parameters used as constraints in some static tools/approaches to model the co-culture of C. autoethanogenum and C. kluyveri.** These values shown are used only when the specific parameter was considered as an input parameter of that specific tool/approach (see S2 Table).

| Parameter | *C. autoethanogenum* | | *C. kluyveri* | Community | |
| --- | --- | --- | --- | --- | --- |
| GEM | iCLAU786 [67] | | ickl708 [68] | | Multi-species GEM [54] |
| CO uptake rate (mmol L^-1^ h^-1^) |  |  | | | 4.8 [53] |
| Relative abundance | 0.4 [54] | | 0.6 [54] | |  |
| Total biomass (g) |  |  | | | 0.22 [53] |
| Growth rate (h^-1^) | 0.021 [53]  min. 0.005 | 0.021 [53]  min. 0.005 | | |  |

**Table S5. Summarized comparison of the available dynamic tools/approaches.**

| **Modeling Tool (Year Developed)** | **Website/GitHub Link** | **Optimization Routine (Single or Bilevel)** | **Programming Language** | **Environment Dependencies** | **Optimization Dependencies** | **Namespace Requirements** | **# of citations (as of December 2022)** |
| --- | --- | --- | --- | --- | --- | --- | --- |
| **DyMMM (2011)** | https://sourceforge.net/p/dymmm/wiki/Home/ | Single | MATLAB | COBRA Toolbox | CPLEX, Mosek, or Gurobi | No | 206 |
| **DFBAlab (2014)** | https://yoric.mit.edu/software/dfbalab/how-obtain-dfbalab#gsc.tab=0 | Single | MATLAB | COBRA Toolbox | CPLEX, Mosek, or Gurobi | No | 83 |
| **MMODES (2019)** | https://mmodes.readthedocs.io/en/latest/ | Bilevel | Python | COBRApy | GLPK | BiGG | 2 |
| **µBialSim (2020)** | https://www.frontiersin.org/articles/10.3389/fbioe.2020.00574/full | Single | MATLAB | CobraToolbox/CellNetAnalyzer | GLPK | No | 27 |

**Table** **S6**. **Inputs, outputs, and assumptions of the dynamic tools/approaches.**

| Tool/approach | Inputs | Outputs | Assumptions |
| --- | --- | --- | --- |
| DyMMM | Genome-scale metabolic models; Initial volume, biomass, and substrate concentrations; duration (time span) of the experimental run; exchange metabolites (reactions) of interest, Michaelis-Menten kinetics | Biomass, substate, and product formation over time, as well as metabolic fluxes at each time step | Reaction rates are constant over time intervals |
| DFBAlab | Genome-scale metabolic models; Initial volume, biomass, and substrate concentrations; duration (time span) of the experimental run; exchange metabolites (reactions) of interest, Michaelis-Menten kinetics | Biomass, substate, and product formation over time as well as metabolic fluxes at each time step | Reaction rates are constant over time intervals |
| MMODES | Genome-scale metabolic models; Initial volume, biomass, and substrate concentrations; duration (time span) of the experimental run; exchange metabolites (reactions) of interest, Michaelis-Menten kinetics | Biomass, substate, and product formation over time as well as metabolic fluxes at each time step | Reaction rates are constant over time intervals |
| µBialSim | Genome-scale metabolic models; Initial volume, biomass, and substrate concentrations; duration (time span) of the experimental run; exchange metabolites (reactions) of interest, Michaelis-Menten kinetics | Two files are generated at the end of the simulation with a date and time stamp in the filename indicating the start of the simulation. The field’s time, compounds, biomass, and mu hold the time, compound concentrations, biomass concentrations, and specific growth rates for each integration step. The field FBA stores data for each FBA model, including the temporal dynamics of all metabolic fluxes, and the mass balance for all exchange reactions. | Reaction rates are constant over time intervals. |

**Table S7. Genome-scale metabolic models (GEMs), GEM constraints, substrate uptake parameters, and initial biomass concentrations used by some dynamic tools/approaches to** **model the co-culture of S. cerevisiae and E. coli** These values shown are used only when the specific parameter was considered as an input parameter of that specific tool/approach (see S2 Table).

| **Parameter** | ***E. coli*** | ***S. cerevisiae*** |
| --- | --- | --- |
| **GEM** | iJR904 | iND750 |
| **GEM Modifications** | Glucose Kinase, GLUK, LB=UB=0  Glucose Exchange, EX_glc(e), LB=UB=0 | - |
| **vgmax (mmol/gDW/h)** | - | 25.9 |
| **Kg (g/L)** | - | 0.5 |
| **vz,max (mmol/gDW/h)** | 9 | - |
| **Kz (g/L)** | 0.01 | - |
| **Ki,e (g/L)** | 8 | 10 |
| **vo,max (mmol/gDW/h)** | 8 | 1.5 |
| **Ko (mmol/L)** | 0.001 | 0.003 |
| **Initial Biomass (gDW/L)** | 0.05 | 0.05 |

**Table S8 Initial substrate concentrations used for the co-culture dynamic tools/approaches model the co-culture of S. cerevisiae and E. coli.**

| **Metabolite** | **Initial Concentration (mmol/L)** |
| --- | --- |
| Glucose | 88.80 |
| Xylose | 53.28 |
| Ethanol | 0 |
| Oxygen | 0.24 |

**Table S9. Summarized comparison of the available spatiotemporal tools/approaches.**

| **Modeling Tool (Year Developed)** | **Website/GitHub Link** | **Optimization Routine (Single or Bilevel)** | **Programming Language** | **Environment Dependencies** | **Optimization Dependencies** | **Namespace Requirements** | **# of citations (as of December 2022)** |
| --- | --- | --- | --- | --- | --- | --- | --- |
| **COMETS (2014, 2021)** | <https://www.runcomets.org> | Single | MATLAB, Python, Command-line | COBRA Toolbox, COBRAPy | CPLEX, Mosek, or Gurobi | BiGG | 312 |
| **BacArena (2017)** | <https://bacarena.github.io> | Single | R | Sybil | CPLEX, Mosek, or Gurobi | No | 144 |
| **IndiMesh (2019)** | <https://journals.plos.org/ploscompbiol/article?id=10.1371/journal.pcbi.1007127> | Bilevel | MATLAB | - | GLPK | No | 32 |
| **CROMICS (2021)** | <https://github.com/EPFL-LCSB/cromics> | Single | MATLAB | - | GLPK | No | 3 |

**Table** **S10**. **Inputs, outputs, and assumptions of the spatiotemporal tools/approaches.**

| Tool/approach | Inputs | Outputs | Assumptions |
| --- | --- | --- | --- |
| COMETS | Genome-scale metabolic models; Initial volume, biomass, and substrate concentrations; duration (time span) of experimental run; exchange metabolites (reactions) of interest, Michaelis-Menten kinetics, Diffusion parameters, Grid size | Biomass, substate, and product formation over time as well as metabolic fluxes at each time step, spatial locations of microbial species | Reaction rates are constant over time intervals. Metabolite diffusion  coefficient is constant |
| BacArena | Genome-scale metabolic models; Initial volume, biomass, and substrate concentrations; duration (time span) of experimental run; exchange metabolites (reactions) of interest, Michaelis-Menten kinetics, Diffusion parameters, Grid size | Biomass, substate, and product formation over time as well as metabolic fluxes at each time step, spatial locations of microbial species | Reaction rates are constant over time intervals. Metabolite diffusion  coefficient is constant. |
| IndiMesh | Genome-scale metabolic models; Initial volume, biomass, and substrate concentrations; duration (time span) of experimental run; exchange metabolites (reactions) of interest, Michaelis-Menten kinetics, Diffusion parameters, Pore size | Biomass, substate, and product formation over time as well as metabolic fluxes at each time step, spatial locations of microbial species | Reaction rates are constant over time intervals. Metabolite diffusion  coefficient is constant |
| CROMICS | Genome-scale metabolic models; Initial volume, biomass, and substrate concentrations; duration (time span) of experimental run; exchange metabolites (reactions) of interest, Michaelis-Menten kinetics, Diffusion parameters, Box size, the volume occupied by cells | Biomass, substate, and product formation over time as well as metabolic fluxes at each time step, spatial locations of microbial species, crowding conditions of the cells | Reaction rates are constant over time intervals. |

**Table S11**. **Genome-scale metabolic models (GEMs) and input parameters used as constraints in some spatiotemporal tools/approaches to model the co-culture of S. enterica and E. coli.** These values shown are used only when the specific parameter was considered as an input parameter of that specific tool/approach (see S2 Table).

| **Parameter** | ***E. coli*** | ***S. enterica*** |
| --- | --- | --- |
| **GEM** | iJO1366 | iiRR1083 |
| **GEM Modifications** | Cystathionine γ-synthase)., *metB*, LB=UB=0 | gain-of-function  mutations in metA (homoserine transsuccinylase)  biomass = S0.reactions.BIOMASS_iRR1083_metals.add_metabolites({met_c : -.5, met_e : .5})  - |
| **Vmax (mmol/gDW/h)** | 10 - | 10 |
| **Km (µM)** | 10 | 10 |
| **Death rate** | 1% | 1% |
| **Metabolite diffusion (cm2/s)** | 0.01 | 0.01 |
| **Biomass diffusion (cm^2^/s)** | 8 | 8 |
| **Max. colony height (200 µm)** | 8 | 8 |
| **Oxygen concentration (µmol/cm^2^)** | 250 | 250 |
