## Supplementary File1 for "A structured evaluation of genome-scale constraint-based modeling tools for microbial consortia"

**Below, we give a brief description of the set of tools and approaches evaluated in this study, based on their usability to model microbial consortia using GEMs.**

**Steady-state methods**

Joint-FBA (2007)

Joint-Flux Balance Analysis (Joint-FBA) was the first approach implemented to model microbial communities, and it is a direct extension of FBA \cite{https://doi.org/10.1038/msb4100131}. Joint-FBA integrates GEM of individual species into a multi-compartment model in which the exchange of metabolites between species occurs through a community compartment. Extracellular fluxes are computed, maximizing the community biomass reaction, defined as the sum of the growth rates of individual species. The stoichiometry of the reaction integrates the species ratio but not as microbial abundances since the extracellular fluxes are not normalized by the total community biomass.

OptCom (2012)

OptCom is a multi-level and multi-objective optimization framework accessed through GAMS for the analysis of microbial communities \cite{Zomorrodi2012}. OptCom predicts relative abundance and flux distributions considering species-level growth rates as the inner problems while relying on community-level objective maximization as the outer problem, for instance, the overall community biomass.

cFBA (2013)

Community Flux Balance Analysis (cFBA) is a translation of FBA applied for single organisms to microbial communities. It is a computational approach that considers the balanced growth of microbial species \cite{Khandelwal2013}, as it occurs in a chemostat. FBA predicts a flux distribution compatible with the optimal value of the objective function and species' relative abundance, maximizing community growth rate as the single objective.

CASINO (2015)

Community And Systems-level INteractive Optimization is a toolbox used to analyze related dietary intake within the human gut microbiota \cite{Shoaie2015}. CASINO performs iterative multi-level optimization to calculate the relative uptake of substrates and metabolite production by each species until the total community biomass production is optimized. The relative abundances are used to constrain the solution space.

SteadyCom (2017)

SteadyCom is a tool integrated with COBRA Toolbox \cite{10.1371/journal.pcbi.1003424}, written in Matlab that predicts microbial composition in a given environment \cite{Hung2017}. SteadyCom assumes that the community grows at a time-averaged constant growth rate, and thus, the organisms grow at the same growth rate. The maximization of the community growth rate becomes a linear problem (LP) once the growth rate is fixed. This allows for the obtaining of a flux distribution and relative abundances for the specified uptake rate.

Microbiome modelling Toolbox (v1, 2018; v2 2022)

Microbiome modelling toolbox (MMT)\cite{Baldini,Heinken} is a tool integrated with COBRA Toolbox, written in Matlab to model microbe–microbe, and host–microbe metabolic interactions within the gut microbiota using GEMs. MMT allows for integrating metagenomic data and human microbial GEMs from AGORA, a computational resource of human microbial GEMs \cite{Heinken2020.11.09.375451}. %MMT informs on pairwise microbe-microbe and host-microbe interactions, and allows for the construction of community models.

Specified communities can be modeled by integrating diets and relative abundance data, and flux distribution can be analyzed by classical multidimensional scaling analyses (PCoA). PCoA produces a Euclidean representation of a group of objects whose association is quantified by any dissimilarity index.

RedCom (2019)

MICOM (2020)

MICOM (\textit{mi}crobial \textit{com}munities) \cite{Diener2020} is a tool implemented as an open-source software package -MICOM- in python. MICOM provides a mechanistic hypothesis that can guide experimental implementations for the gut microbiota. It predicts community, species growth rates, and metabolic fluxes, allowing the incorporation of relative abundance and AGORA models.

MICOM integrates two optimization steps: i) the community growth rate is fixed to a fraction of its optimum value, and ii) the L2 -regularization function- norm of individual growth rates is minimized. This optimization strategy is motivated by the assumption that there is a trade-off between community growth rate and individual taxon growth rates, since most microbial communities have many species with small but nonzero abundances.

NECom (2021)

Nash Equilibrium predictor for microbial Communities -NECom- follows a bi-level approach that does not consider 'force-altruism' to study microbial interactions \cite{Cai2021}, but Nash Equilibria (NE). A microbial community is in a NE if the strategy of each microbe is maximizing its growth rate simultaneously such that no species can increase its performance by changing its strategy unilaterally. In NECom, any feasible solution is NE of community models with or without an outer-level objective or imposed strategy. NECom implements standard FBA to maximize biomass production as the inner problem satisfying substrate availability, thermodynamics, and mass balance principles.

**Dynamic methods**

DyMMM (2011)

Dynamic Multispecies Metabolic Modeling (DyMMM) \cite{Zhuang2011} was the first dynamic framework developed to examine microbial communities at the genome scale. DyMMM optimizes specific growth rates for each species via maximizing the biomass function of each species. Subsequently, the predicted exchange fluxes are used to update the extracellular environment. Then, time-based changes in biomass and metabolite concentrations can be determined, and corresponding consumption and production fluxes can be estimated by using Michaelis-Menten kinetic expressions.

DFBALab (2014)

Dynamic Flux Balance Analysis laboratory (DFBALab) \cite{Gomez2014} is an approach similar to DyMMM, which seeks to alleviate infeasible FBA problems and degeneracy associated with shared metabolites from uptake and secretion fluxes. This fix is achieved by arranging the objectives according to priority. In particular, the first objective is biomass maximization, and the following maximization/minimization is performed on specific metabolites from exchange fluxes. DFBALab requires a user-defined list of shared metabolites to avoid improbable solutions.

d-OptCom (2014)

Dynamic-OptCom (d-OptCom) is an extension of the static method OptCom \cite{Zomorrodi2014}. It utilizes temporal measurements to explore how the structure of a microbial community develops over time. More specifically, d-OptCom uses a multi-objective, bilevel framework present in OptCom, which incorporates a total community biomass concentration maximization as an outer-level objective while maintaining mass conservation to biomass constraints to the outer problem.

MCM (2015)

Microbial Community Modeler (MCM) \cite{Louca2015} is a tool that simulates dynamic community models using dFBA. The tool’s added features are that it can perform construction, calibration, and statistical analysis of microbial community models based on experimental data. MCM was initially devised for a homogeneous \textit{E. coli} community and later applied to investigate species assemblages in nitrifying and methanogenic bioreactors \cite{louca2017taxonomic}.

DAPHNE (2019)

DynAmic Population HeterogeNEity (DAPHNE) \cite{Succurro2019} is a dynamic modeling framework that embeds FBA within differential equations. In particular, this approach builds on previous dFBA methods by employing the minimization of metabolic adjustment (MOMA) algorithm \cite{segre2002analysis} in dFBA to minimize the metabolic adjustments between various time points. This tool also incorporates certain mechanisms into the framework, such as population transitions not accounted for in previous approaches.

μbialSim (2020)

µbialSim \cite{Popp2020} is a recent dFBA-based modeling tool that can predict dynamic shifts in the environment and composition of a microbial community that contains several species either in batch or chemostat mode. Each species can share a supply of metabolites that can be exchanged among the species. It also uses an augmented Euler method to prevent negative metabolite concentrations. This implementation potentially decreases simulation times as well. FBA simulations in µbialSim can be performed using functions from either COBRA Toolbox \cite{heirendt2019creation} or CellNetAnalyzer \cite{klamt2007structural}. µbialSim has been used to study ranging systems from syntrophic methanogenic co-culture to a 773-species human gut microbiome \cite{Popp2020}.

MMODES (2020)

Metabolic Models based Ordinary Differential Equations Simulation (MMODES) \cite{Garcia-Jimenez2020} is a tool that combines GEMs and ODEs to temporally simulate biomass and metabolite dynamics. This framework was greatly influenced by DAPHNE. However, this method was developed for general cases. MMODES also can include perturbations via changes in the medium or adjusting specific biomass or metabolite concentrations of certain species.

SurfinFBA (2020)

SurfinFBA \cite{10.1371/journal.pcbi.1007786} is a GEM modeling tool that allows for the efficient dynamic modeling of microbial communities. This increase in efficiency compared to previous tools is leveraged by selecting a flux vector basis for the space of internal fluxes for each microbe in a community. Hence, this basis is used to simulate forward by solving a relatively inexpensive system of linear equations at most time steps. Also, as the solution becomes infeasible, surfinFBA permits a different but related optimization problem to form a reasonable basis to continue the forward simulation. Overall, the surfinFBA algorithm was shown to use fewer optimizations than other dFBA approaches.

**Spatiotemporal methods**

MatNet (2013)

MATLAB-NetLogo (MatNet) \cite{10.1371/journal.pone.0078011} modeling framework extension was the first multiscale modeling tool published for spatiotemporal genome-scale metabolic modeling. The tool incorporates both an individual-based model (IBM) and constraint-based metabolic modeling. This approach using IBMs allows for studying how interaction mechanisms lead to the formation of structures or patterns. However, IBM simulations are computationally expensive and increase considerably depending on model complexity. MatNet connected MATLAB with the NetLogo platform to manage tractability issues and was able to evaluate \textit{Pseudomonas aeruginosa} biofilm formation\cite{10.1371/journal.pone.0078011}.

COMETS (v1, 2014; v2, 2021)

Computation of Microbial Ecosystems in Time and Space (COMETS) \cite{Harcombe2014,Dukovski2021} was the first widely available tool for the spatiotemporal modeling of microbial communities. COMETS represents cellular biomass on a population level where cells are simulated per grid position. A recent version of this framework (COMETS 2) \cite{Dukovski2021} features more accurate biophysical models that include enzyme functionality and account for evolutionary dynamics. COMETS is a flexible tool compatible with the most widespread interfaces, such as MATLAB, Python, and command-line, and even supports a graphical user interface (GUI). However, COMETS has some difficulty in terms of scalability when solving complex convection-diffusion equations.

BacArena (2017)

BacArena \cite{Bauer2017} is a tool that couples constraint-based GEM metabolic modeling with IBM. It is a framework that is beneficial when simulating heterogeneous cell populations. Each GEM is allocated an individualized two-dimensional grid where metabolic exchange occurs. Then, the individual and environmental metabolite concentrations are updated at each time step. Another feature of BacArena is that it can be used to model cross-feeding mechanisms between species.

IndiMeSH (2019)

Individual bacterial agents in heterogeneous and dynamic soil habitats (IndiMeSH) is a novel modeling framework that uses angular pore networks to represent a microbial habitat coupled with IBM to characterize bacterial cells and GEMs to define the metabolic networks \cite{10.1371/journal.pcbi.1007127}. Furthermore, this framework realistically simulates mass transfer limitations in the gas and liquid phase. Therefore, this approach can enable predictions of specialized nutrient diffusion settings.

MiMoSA (2019)

Multiscale Multiobjective Systems Analysis (MiMoSA) \cite{Gardner2019} is a metabolic modeling tool that can simulate individual cells spatiotemporally, model the diffusion of nutrients and light, and depict the different cell interactions with each other and the environment. This approach involves inputting both continuous and discrete variables as well as various expressions formalisms to represent the multilevel behavior in cellular communities. Hence, an IBM framework was incorporated into this approach to reflect the direct interaction of different levels throughout metabolism, environment, and cellular space.

ACBM (2020)

Agent and constraint-based modeling (ACBM) \cite{Karimian2020} framework is an approach that simulates a cell population’s spatiotemporal dynamics and interactions in three-dimensional space. ACBM employs an IBM conceptualization of biomass. Therefore, this tool is limited in its scalability to smaller communities. Also, ACBM contains a user-friendly GUI panel to increase the accessibility of the tool to novices.

CROMICS (2021)

CROwding-Modeling of In-silico Community System (CROMICS) \cite{Angeles-Martinez} is an approach that combines techniques such as IBM and thermodynamic flux analysis (TFA) to detect the metabolic variability within a microbial community. For estimating the metabolic activity of individual cells under local environmental conditions, CROMICS uses GEMs. In addition, this tool models a crowding effect within the volume among the cells by using scaled particle theory (SPT) \cite{Lebowitz1965}.
